# A dead gene walking: convergent degeneration of a clade of MADS-box genes in Brassicaceae

**DOI:** 10.1101/149484

**Authors:** Andrea Hoffmeier, Lydia Gramzow, Amey S. Bhide, Nina Kottenhagen, Andreas Greifenstein, Olesia Schubert, Klaus Mummenhoff, Annette Becker, Günter Theißen

## Abstract

Genes are ‘born’, and eventually they ‘die’. In contrast to gene birth, however, gene death has found only limited scientific interest, even though it is of considerable evolutionary importance. Here we use B_sister_ genes, a subfamily of MIKC-type MADS-box genes, as a model to investigate gene death in unprecedented detail. Typical MIKC-type genes encode conserved transcription factors controlling plant development. We show that *ABS*-like genes, a clade of B_sister_ genes, are indeed highly conserved in Brassicaceae maintaining the ancestral function of B_sister_ genes in ovule and seed development. In contrast, their closest paralogs, the *GOA*-like genes, have been undergoing convergent gene death in Brassicaceae. Intriguingly, erosion of *GOA*-like genes occurred after millions of years of co-existence with *ABS*-like genes. We thus describe Delayed Convergent Asymmetric Degeneration (DCAD), a so far neglected but possibly frequent pattern of duplicate gene evolution that does not fit classical scenarios. DCAD of *GOA*-like genes may have been initiated by a reduction in the expression of an ancestral *GOA*-like gene in the stem group of Brassicaceae and driven by dosage subfunctionalization. Our findings have profound implications for gene annotations in genomics, interpreting patterns of gene evolution and using genes in phylogeny reconstructions of species.

## INTRODUCTION

The origin (‘birth’) and loss (‘death’) of genes are considered to be of major importance for the phenotypic evolution of organisms (Flagel and Wendel, 2009; Smith and Rausher, 2011). Understanding the dynamics and interdependence of these processes is thus of utmost biological interest. Many genes originate by duplication of already existing genes. The fate of such duplicate genes has been a subject of research for decades and different evolutionary trajectories of duplicate genes have been both hypothesized and observed (Ohno, 1970; Ohta, 1987; Lynch and Conery, 2000).

Duplicated genes are often assumed to be initially redundant (Lynch and Conery, 2000; Lynch and Force, 2000). Since deleterious mutations are much more frequent than beneficial ones (Charlesworth and Charlesworth, 1998; Keightley and Lynch, 2003), the most likely fate is that one of the pair of duplicated genes will degenerate into a pseudogene, a process termed ‘nonfunctionalization’, while the other one may be maintained under purifying selection for the ancestral function (Ohno, 1970). The pseudogene will eventually be lost from the genome (Prince and Pickett, 2002). However, it is also possible that both copies acquire loss-of-function mutations after the gene duplication. If those mutations are complementary and affect independent sub-functions, both genes together are required to fulfill the full complement of functions of the single ancestral gene (Force et al., 1999; Prince and Pickett, 2002). This process is termed ‘subfunctionalization’. Finally, a third classical fate of duplicate genes, termed ‘neofunctionalization’, has been considered. It occurs when after gene duplication one of two copies undergoes directional selection to perform a novel function while the other copy maintains the ancestral gene function (Ohno, 1970; Force et al., 1999). Neofunctionalization is the only fate of the three ones mentioned that leads to genes with truly new functions and can thus explain how biological novelties evolve (Francino, 2005; He and Zhang, 2005).

Since the different possible fates, especially sub- and neofunctionalization, can also affect the same genes (resulting in ‘subneofunctionalization’; (He and Zhang, 2005)), the long-term evolution of duplicate genes is difficult to understand and has not yet been studied intensively in plants. A comprehensive understanding requires detailed comparative studies of suitable model genes and species. Here, we introduce the clades of *ABS*- and *GOA*-like genes from the Brassicaceae for such an analysis. The two gene clades have been named after the two B_sister_ genes of *Arabidopsis thaliana*, namely *ARABIDOPSIS BSISTER* (*ABS*, also known as *TT16* and *AGL32*) (Becker et al., 2002; Nesi et al., 2002) and *GORDITA* (*GOA*, also known as *AGL63*) (Erdmann et al., 2010; Prasad et al., 2010). B_sister_ genes belong to the MIKC-type MADS-box genes encoding transcription factors. MIKC-type transcription factors control essential ontogenetic processes in plants, such as flower and fruit development (for a review see Gramzow and Theißen (2010); Smaczniak et al. (2012)). MIKC-type proteins exhibit a conserved domain structure, where the MADS (M) domain is followed by an Intervening (I), a Keratin-like (K), and a C-terminal (C) domain (Ma et al., 1991). At least some MIKC-type proteins have the capacity to form multimeric transcription factor complexes in a combinatorial manner (Honma and Goto, 2001; Theißen and Saedler, 2001; Smaczniak et al., 2012; Theißen et al., 2016). Like many genes encoding transcription factors, MIKC-type genes have been preferentially retained after whole genome duplications (WGD) and have been conserved quite well during flowering plant evolution (Shan et al., 2009; Gramzow and Theißen, 2010).

Typically, B_sister_ genes have important functions in the development of female reproductive organs and seeds (Becker et al., 2002; Nesi et al., 2002; Mizzotti et al., 2012; Yang et al., 2012; Xu et al., 2016). This may well represent the ancestral role of these genes, which possibly had already been established in the most recent common ancestor of extant seed plants (Becker et al., 2002). In this study, we analyze the birth and fate of the clades of *ABS*- and *GOA*-like genes resulting from a duplication or triplication of an ancestral B_sister_ gene near the base of core eudicots. Our data reveal in detail how a complete clade of ancient orthologs was predisposed to extinction in the plant family of Brassicaceae by convergent non-functionalization and then eventually lost many of its members over an extended period of time. We show that in such cases comprehensive phylogenetic analyses have the potential to identify putative pseudogenes even if the respective sequences are still lacking unequivocal hallmarks of pseudogenization. Our findings have profound consequences for the annotation of genes in genomics projects, interpreting molecular patterns of gene evolution and for using genes in phylogeny reconstructions of species.

## RESULTS

### Origin of *ABS*- and *GOA*-like genes predates the origin of Brassicaceae

To clarify the origin of ABS- and GOA-like genes we searched for Bsister genes in whole genomes of species at informative positions of the angiosperm phylogeny. We identified 78 B_sister_ genes from 51 plant species. Furthermore, we sequenced the genomic region of the two B_sister_ genes from *Lepidium campestre* and cDNA of nine B_sister_ genes from six other species (Supplemental Data Set 1). Using these data and published B_sister_ gene sequences (Leseberg et al., 2006; Díaz-Riquelme et al., 2009; Erdmann et al., 2010; Yang et al., 2012; Chen et al., 2013), we reconstructed phylogenetic trees based on two different methods, Bayesian inference (BI) and Maximum Likelihood (ML). The robustness of our results is supported by the similarity of the topology of the different trees (Figure 1, Supplemental Figures 1 and 2). In our phylogenies, B_sister_ genes from gymnosperms, monocots, and eudicots form separate clades, thus reflecting speciation events. For core eudicot B_sister_ genes, our phylogenetic trees suggest the existence of two large clades with high support values. One of these putative clades contains *ABS* and the other one includes *GOA* of *A. thaliana*. Hence, we termed these clades *ABS*- and *GOA*-like genes, respectively. The clade of *GOA*-like genes comprises genes from a number of rosid and asterid species while the clade of *ABS*-like genes includes genes from a number of rosids but is lacking asterid species. The B_sister_ genes identified in the basal eudicot species *Aquilegia coerulea*, *Nelumbo nucifera* and *Eschscholzia californica* branch off before the duplication of *ABS*- and *GOA*-like genes. These findings indicate that the duplication giving rise to *ABS*- and *GOA*-like genes likely occurred after basal eudicots branched off but before the radiation of core eudicots, and thus long before the origin of Brassicaceae. Hence, our data are compatible with the hypothesis that this gene duplication occurred during the γ-whole genome triplication about 120 million years ago (MYA) in a common ancestor of asterids and rosids (Jiao et al., 2012). During the γ-genome triplication, a third B_sister_ gene should have originated as well. However, since we only observe two large clades of B_sister_ genes with our sampling, this third gene may have been lost quickly.

**Figure 1:**
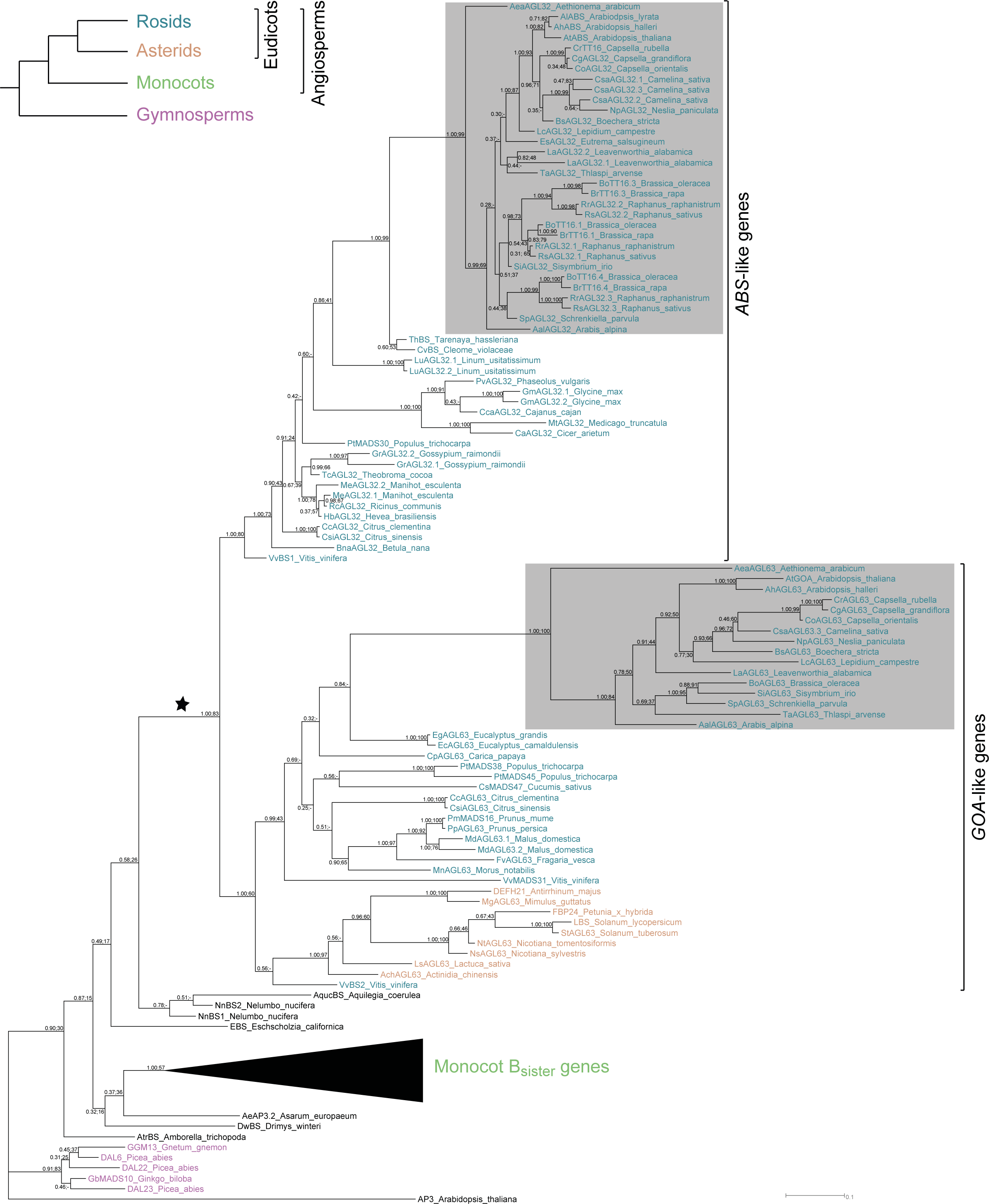
Phylogeny of _B_sister__ genes of seed plants as reconstructed by Bayesian inference based on protein sequences. The duplication event of the _B_sister__ genes resulting in *ABS*- and *GOA*-like genes is indicated by a star and the clades of *ABS*- and *GOA*-like genes are marked on the right. The *ABS*- and *GOA*-like genes of Brassicaceae are indicated by shading. Branch lengths are drawn proportional to the number of substitutions. The left value at each node represents the posterior probability of the corresponding clade as calculated by MrBayes, while the right value indicates the bootstrap value of the Maximum Likelihood phylogeny (Supplemental Fig. 2); if the corresponding clade did not occur in the Maximum Likelihood phylogeny a ‘-‘ is shown. *APETALA3* (*AP3*) was used as representative of the outgroup. The fully resolved phylogeny is shown in Supplemental Figure 1. In the upper left corner, a simplified phylogeny of seed plants is given with a color code which we used for the large gene phylogeny.

We then investigated the gene order surrounding *ABS*- and *GOA*-like genes in core eudicots to corroborate the independence of the two gene clades (Figure 2). We detected that a conserved synteny of *ABS*-like genes can be traced from Brassicaceae to the early diverging rosid species *Vitis vinifera* (Figure 2A). This confirms the finding of our phylogeny that these genes form a major gene clade. For the clade of *GOA*-like genes, a conserved synteny from Brassicaceae to *V. vinifera* was observed as well (Figure 2B). However, the genomic organization of the investigated asterid species showed that none of the genes flanking the B_sister_ gene is orthologous to the genes flanking the *GOA*-like gene in rosid species. In general, none of the annotated genes surrounding *GOA*-like genes appears to be orthologous to any annotated gene surrounding *ABS*-like genes and *vice versa*. These findings support the data of the phylogeny reconstruction showing that two major clades of B_sister_ genes, *ABS*- and *GOA*-like genes, are present in rosids. We also searched for orthologous genes surrounding the B_sister_ genes in the basal eudicots *N. nucifera* and *A. coerulea* but were unable to find orthology to the surrounding genes of either *ABS-* or *GOA*-like genes.

**Figure 2:**
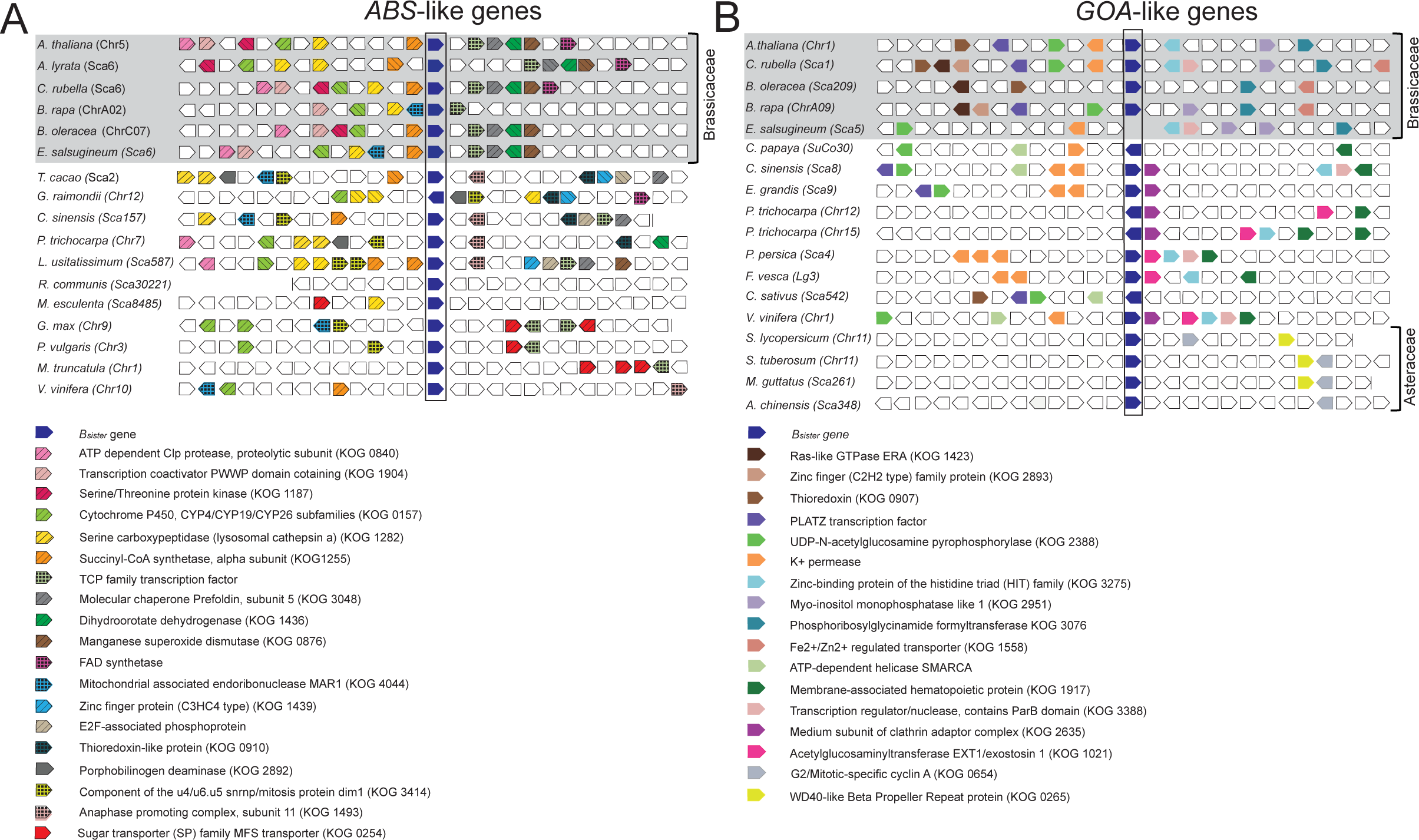
Genomic organization (synteny) of B_sister_ genes in core eudicots. *ABS*- **(A**) and *GOA*-like genes **(B)** are boxed. Genes surrounding B_sister_ genes and their orientation are indicated by pentagons. Putative orthologs among the surrounding genes which are present in more than two species are shown in identical color and pattern. The annotation of these genes according to the euKaryotic Orthologous Groups (KOG, identifier given in brackets) or the function as annotated in Phytozome is given at the bottom. For Brassicaceae species in which more than one *ABS-* or *GOA*-like gene is present, one gene was chosen arbitrarily. The end of a scaffold is marked by a vertical line. The *GOA*-like gene and the neighboring 3’ gene in *E. salsugineum* have been lost. Genomic loci of B_sister_ genes of Brassicaceae species are highlighted in gray. Abbreviations: Chr, chromosome; Sca, scaffold; SuCo, supercontig; Lg, linkage group

### Frequent loss of either *ABS-* or *GOA*-like genes in early eudicot evolution

To clarify the early evolution of Bsister genes in eudicots we mapped the presence of ABS- and *GOA*-like genes onto a phylogeny of extant eudicots with sequenced genomes (Figure 3). All of the analyzed genomes encode at least one B_sister_ gene pointing to the high importance of this gene clade. Moreover, in *V. vinifera*, *Citrus sinensis*, *Citrus clementina*, *Populus trichocarpa* and in most of the Brassicaceae species we identified genes of both clades, *ABS*-and *GOA*-like genes. According to our B_sister_ gene phylogeny and the phylogeny of eudicots, we deduce that both *ABS*- and *GOA*-like genes were lost at least five times independently during the evolution of eudicots outside Brassicaceae (Figure 3). We searched the literature and the NCBI databases for expressed sequence tags (EST) and short read archives (SRA) (Sayers et al., 2012) to find out the expression domains of *ABS*- and *GOA*-like genes in core eudicots. The tissue resolution is quite divergent between different studies (Supplemental Figure 3). Nevertheless, a picture emerges showing expression of both types of B_sister_ genes in flowers, where expression was mainly found in ovules when detailed tissue analyses had been conducted. Furthermore, expression of both genes was often detected in fruits where seeds were mainly identified as the source of expression if studied in more detail. Hence, *ABS*- and *GOA*-like genes have similar expression patterns, indicating little expression divergence between these genes after the duplication event. It is thus conceivable that *ABS*- and *GOA*-like genes may have had a more or less redundant function for a period of up to 120 million years after their duplication.

**Figure 3:**
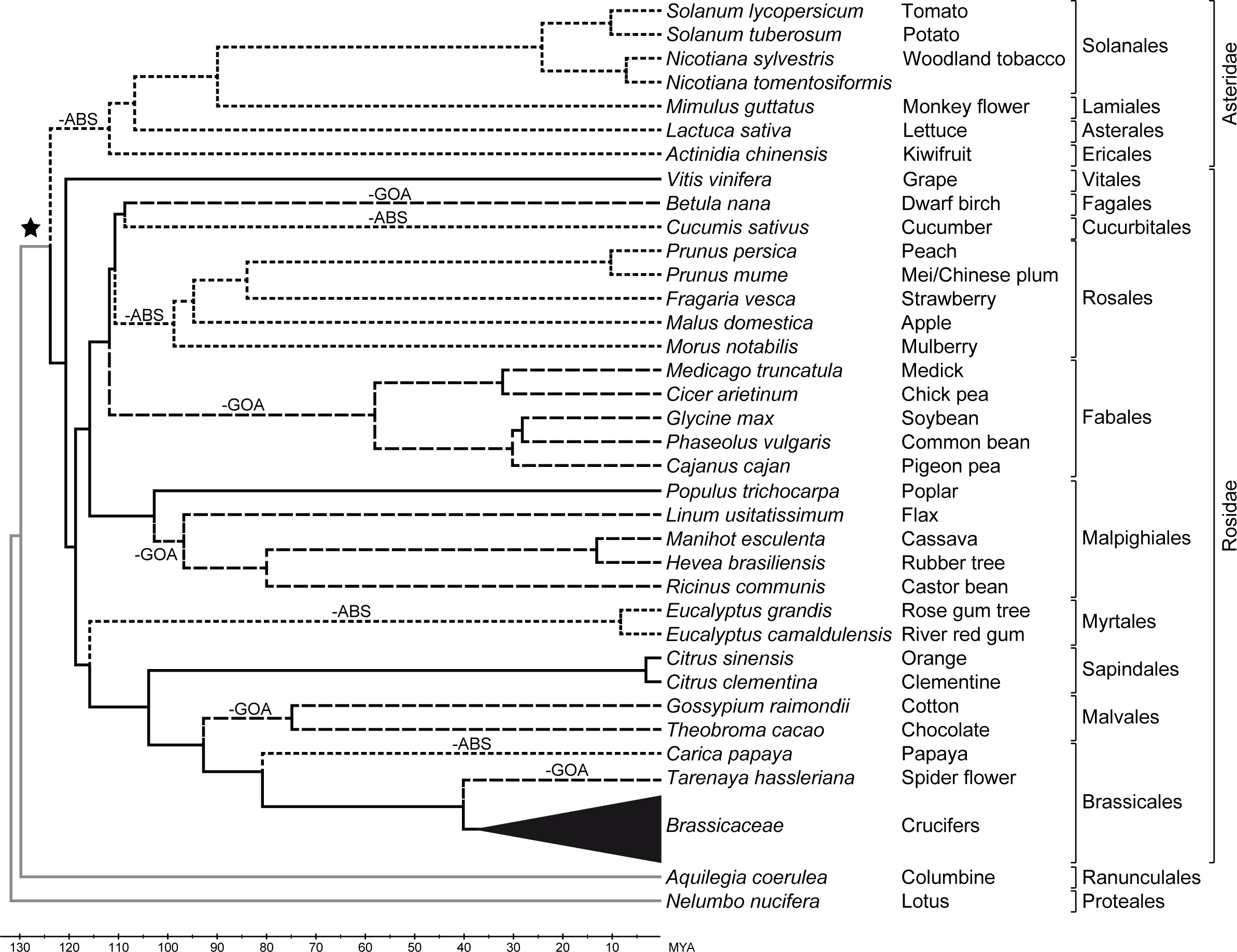
Independent losses of *ABS*- and *GOA*-like genes from genomes of different eudicots. A phylogeny of eudicot species in which B_sister_ genes were analyzed is shown. The phylogeny was drawn according to Magallón et al. (2015) using the estimated divergence times in Million Years Ago (MYA) as given therein. Taxonomic groups are indicated at the right margin. Branches leading to lineages having both *ABS*- and *GOA*-like genes are shown in solid black lines; branches leading to lineages which do not have *GOA*-like genes are shown in broken lines; branches leading to lineages which do not have *ABS*-like genes are shown in dotted lines. Loss of *ABS*- and *GOA*-like genes is marked by –ABS and –GOA, respectively on the corresponding branches. Branches of lineages which diverged before the duplication of *ABS*- and *GOA*-like genes are shown in gray. The stem group in which the duplication event into *ABS*- and *GOA*-like genes likely happened, is indicated by a star. For simplification, the Brassicaceae species are combined into a triangle. A resolved phylogeny for Brassicaceae species can be found in Figure 4.

## Parallel nonfunctionalization of *GOA*-like genes in Brassicaceae

Given the many genomics resources available in Brassicaceae, we decided to study the evolution of *ABS*- and *GOA*-like genes in this plant family in more detail. Our analyses comprise 22 different species and cover the two major Brassicaceae lineages – lineage I and expanded lineage II (Beilstein et al., 2006). We found several species which lack a *GOA*-like gene, as well as frequent losses of *GOA*-like genes after whole genome duplications in the subgenomes of several species (Figure 4). In *Eutrema salsugineum* we found a small scale deletion including the *GOA*-like gene, while the surrounding genes are largely conserved (Figure 2B). In the *GOA*-like gene of *Arabidopsis lyrata*, we identified an insertion of a transposon into exon 3, likely leading to nonfunctionalization. The genera *Brassica* and *Raphanus* share a whole genome triplication event (Lysak et al., 2005; Moghe et al., 2014) leading to potentially three *ABS*-like genes which we identified. However, no *GOA*-like gene was found in the genomes of *R. sativus* and *R. raphanistrum* and only one *GOA*-like gene was observed in the *B. oleracea* and *B. rapa* genomes. Moreover, the *GOA*-like gene in the sequenced genome of *B. rapa* as well as that of several other *B. rapa* subspecies shows a deletion of one nucleotide in the MADS box (Supplemental Figure 4) leading to a frameshift and consequently to a premature stop codon causing pseudogenization. In *Lepidium meyenii* (Zhang et al., 2016), which is a disomic octoploid, we could identify four normal-appearing *ABS*-like genes, as expected. In contrast, we were only able to find the remnants of two *GOA-*like genes with large deletions and early premature stop codons, and no traces of the other two expected *GOA-*like genes.

**Figure 4:**
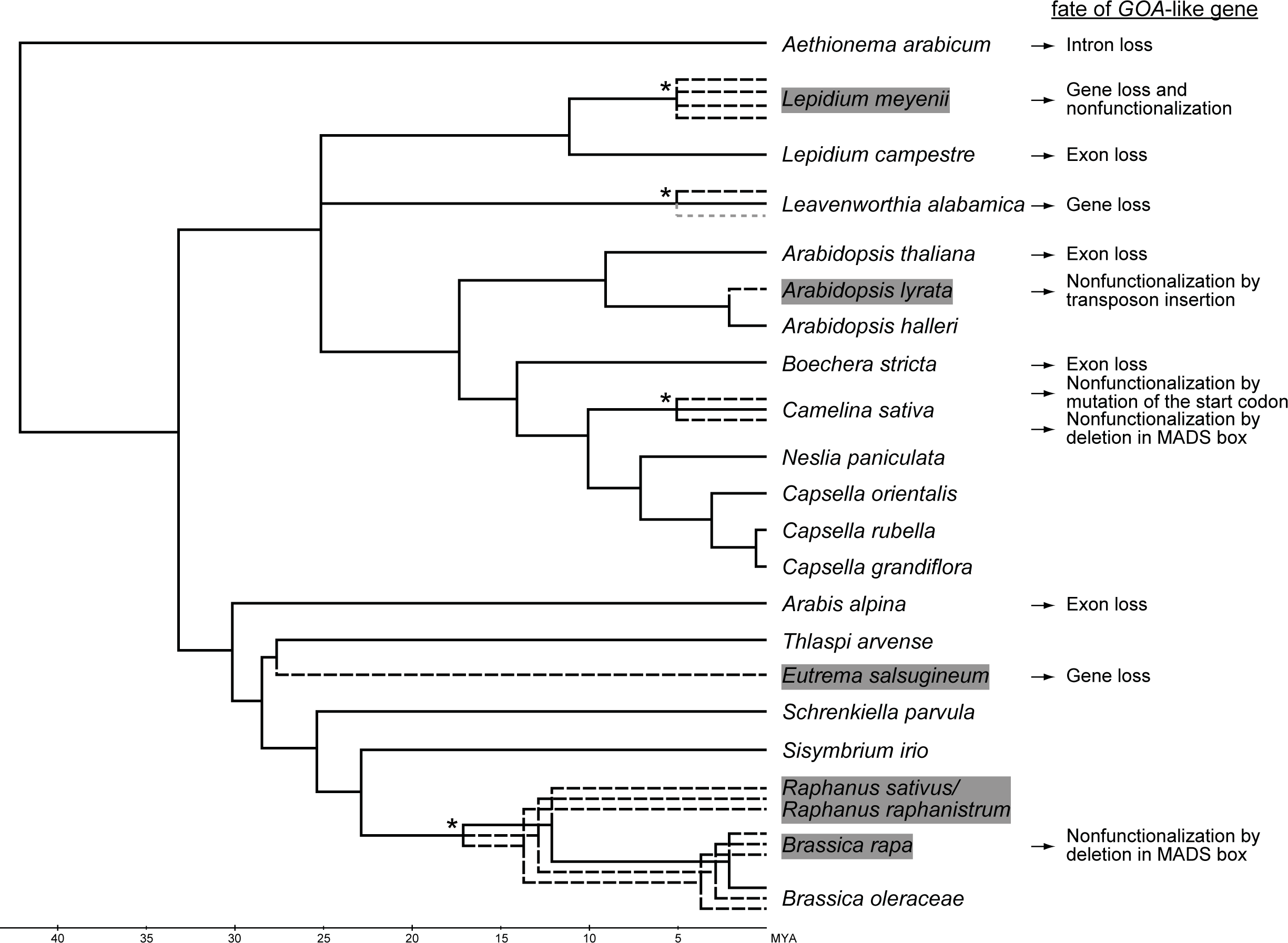
Independent losses of *GOA*-like genes from genomes of Brassicaceae. The species phylogeny is based on publications by Guo et al. (2017) and Huang et al. (2016) using the estimated divergence times in Million Years Ago (MYA) as given therein. The phylogeny contains only species for which whole genome information is available and *L. campestre*. Branches leading to genomes or subgenomes containing both *ABS*- and *GOA*-like genes are shown in solid lines; branches leading to genomes or subgenomes which do not have *GOA*-like genes are shown in broken lines. Six species (*A. lyrata*, *B. rapa, E. salsugineum, L. meyenii, R. sativus* and *R. raphanistrum*) for which we did not find a functional *GOA*-like gene are highlighted in grey. The fate of the corresponding *GOA*-like genes is given on the right. Stars indicate whole genome polyploidization events. The grey dotted branch leading to *L. alabamica* indicates that we were neither able to detect a third syntenic region for *ABS*- nor for *GOA*-like genes.

In *Camelina sativa* a whole genome triplication has occurred (Kagale et al., 2014), but only one, most likely functional *GOA*-like gene could be identified. In the other two loci we found mutations in the start codon and in the middle of the MADS box, respectively, which probably led to nonfunctionalization of these genes. For *Leavenworthia alabamica*, which also underwent a recent whole genome triplication (Haudry et al., 2013), only one *GOA*-like gene could be found. In a second region, which is syntenic to the region of *GOA*, a large number of genes have been lost, including the *GOA*-like gene. Furthermore, we did not succeed in identifying the third syntenic region of *GOA*-like genes.

Taken together, we did not find a potentially functional *GOA*-like gene in the published genome sequences of six species (*A. lyrata*, *B. rapa, E. salsugineum, L. meyenii, R. raphanistrum* and *R. sativus*), whereas not a single Brassicaceae species without a potentially functional *ABS*-like gene in its genome was identified. Including intraspecific paralogs, we noted eleven events of pseudogenization of a *GOA*-like gene, but no such case for an *ABS*-like gene (Figure 4). The mechanisms by which the *GOA*-like genes were pseudogenized as well as the positions in the phylogeny are diverse, indicating independent events. In contrast to the situation in taxa other than Brassicaceae, where *ABS*- and *GOA*-like genes appear to have been lost with similar frequency (Figure 3), our data clearly indicate that *GOA*-like genes were lost preferentially in the analyzed Brassicaceae species (Figure 4).

### In Brassicaceae *GOA*-like genes show less purifying selection than *ABS*-like genes

To find out whether the preferential loss of *GOA*-like genes in Brassicaceae is due to reduced selection pressure, we conducted a selection analysis determining the ratio of the rates of nonsynonymous to synonymous substitutions (ω). We used a dataset including only genes from species which have retained both, *ABS-* and *GOA*-like genes, and the B_sister_ gene *AqucBS* from *A. coerulea* as an outgroup representative (Figure 5). The five ratio model, allowing different ω ratios for *ABS*- and *GOA*-like genes in Brassicaceae (ω1 and ω2), the branches leading to *ABS*- and *GOA*-like genes in Brassicaceae (ω3 and ω4) and all remaining branches (ω0), fitted the data best (Supplemental Table 1). All ω ratios are well below 1, indicating that the coding regions of eudicot B_sister_ genes have been under substantial purifying selection suggesting a functional importance of the encoded gene products at least during longer periods of their evolutionary history. However, the ω ratio for *GOA*-like genes (0.46) is much higher than that for *ABS*-like genes (0.23) in Brassicaceae, indicating that purifying selection was considerably relaxed in the coding regions of *GOA*-like genes during the evolution of Brassicaceae.

**Figure 5:**
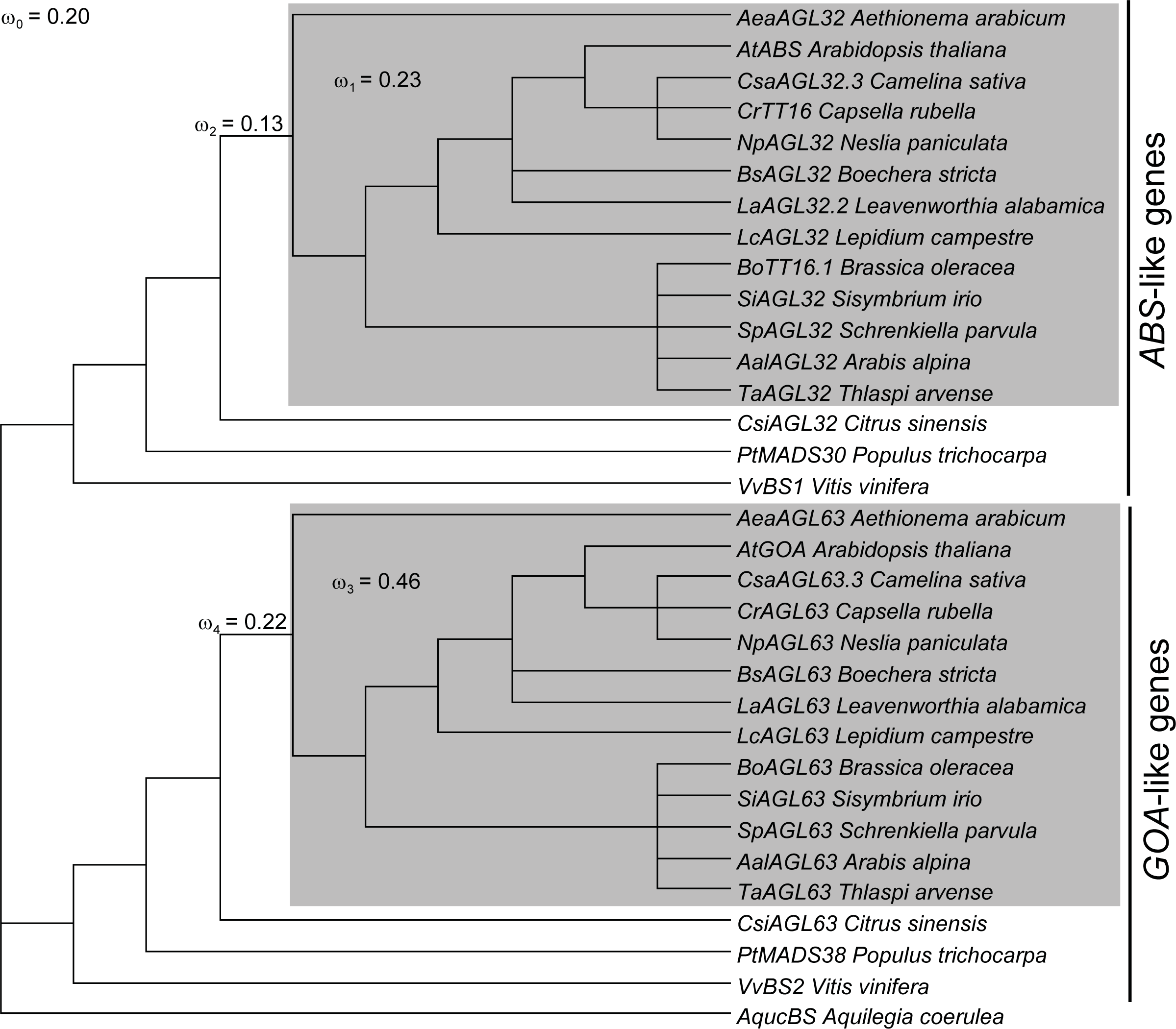
Selection analysis of B_sister_ genes. Ratios of nonsynonymous to synonymous substitution rates (dN/dS = ω) are indicated as calculated according to the given phylogeny and a five ratio model. The phylogeny includes *ABS*- and *GOA*-like genes from eudicots which have retained both types of B_sister_ genes and of the B_sister_ gene of *Aquilegia coerulea*. The phylogeny is based on a species phylogeny from Koenig and Weigel (2015). The clades of *ABS*- and *GOA*-like genes are marked on the right. B_sister_ genes of Brassicaceae are indicated by shading. The five ratio model allows different ω values for all branches of *ABS*-like genes in Brassicaceae (ω_1_), for the branch leading to the *ABS*-like genes in Brassicaceae (ω_2_), for all branches of *GOA*-like genes in Brassicaceae (ω_3_), for the branch leading to the *GOA*-like genes in Brassicaceae (ω_4_) and another value for all remaining branches (ω_0_).

### The exon-intron structure of *ABS*-like genes is more conserved than that of *GOA*-like genes

To analyze the conservation of whole genomic loci, we studied the exon-intron structure and pairwise sequence identities of the exon and intron sequences of *ABS*- and *GOA*-like genes. In Table 1 and in the following, exons are numbered according to their homologous exons in *Thlaspi arvense* (Figure 6). While all *ABS*-like genes are comprised of six coding exons, this number varies between four and six for *GOA*-like genes (Figure 6). Separate alignments of the genomic sequences of *ABS-* and *GOA*-like genes from Brassicaceae showed that exons are generally more conserved than introns, as expected (Table 1; Figure 6). For *ABS*-like genes, the mean value of the pairwise identities between exons is 87% while it is 69% between introns. For *GOA*-like genes, exons are on average 77% identical, while introns are 63% identical. The variable number of coding exons as well as the significantly lower conservation of exons and introns corroborates our view that there has been less selection pressure on *GOA*-like genes than on *ABS*-like genes and reveals that the whole genomic loci of *GOA*-like genes, not just their coding regions, have generally been less conserved than that of *ABS*-like genes in Brassicaceae.

**Table 1:** Average pairwise identities ± SDs between exons and introns of *ABS*- and *GOA*-like genes in Brassicaceae. The highest value for each exon and intron is highlighted in bold. # indicates a significant difference between *ABS*- and *GOA*-like genes (*P ≤* 0.05, two-tailed Mann-Whitney U-test). Exons and introns are numbered according to the *Thlaspi arvense* B_sister_ genes (Figure 6).

|  | Exon 1 | Exon 2 | Exon 3 | Exon 4 | Exon 5 | Exon 6 | Mean value |
| --- | --- | --- | --- | --- | --- | --- | --- |
| *ABS*-like genes | **88.6 % ± 3.8 #** | **84.5 % ± 8.4 #** | **85.4 % ± 6.1 #** | **92.2 % ± 3.5 #** | **89.1 % ± 5.6 #** | **82.4 % ± 5.6 #** | **87.1 % ± 14.1** |
| *GOA*-like genes | 83.5 % ± 6.6 | 80.3 % ± 13.1 | 81.4 % ± 7.5 | 75.4 % ± 15.8 | 70.6 % ± 16.8 | 69 % ± 12.4 | 76.7 % ± 30.9 |
|  | Intron 1 | Intron 2 | Intron 3 | Intron 4 | Intron 5 |  |  |
| *ABS*-like genes | **66.2 % ± 10.3 #** | 61.4 % ± 10.8 | **80.5 % ± 30.2 #** | **67.4 % ± 18.6 #** | **71.2 % ± 22.8 #** | | **69.3 % ± 44.7** |
| *GOA*-like genes | 58.9 % ± 8.5 | **61.8 % ± 9.3** | 72.3 % ± 19.4 | 65.7 % ± 11.9 | 58.4 % ± 10.5 | | 63.4 % ± 28.1 |

**Figure 6:**
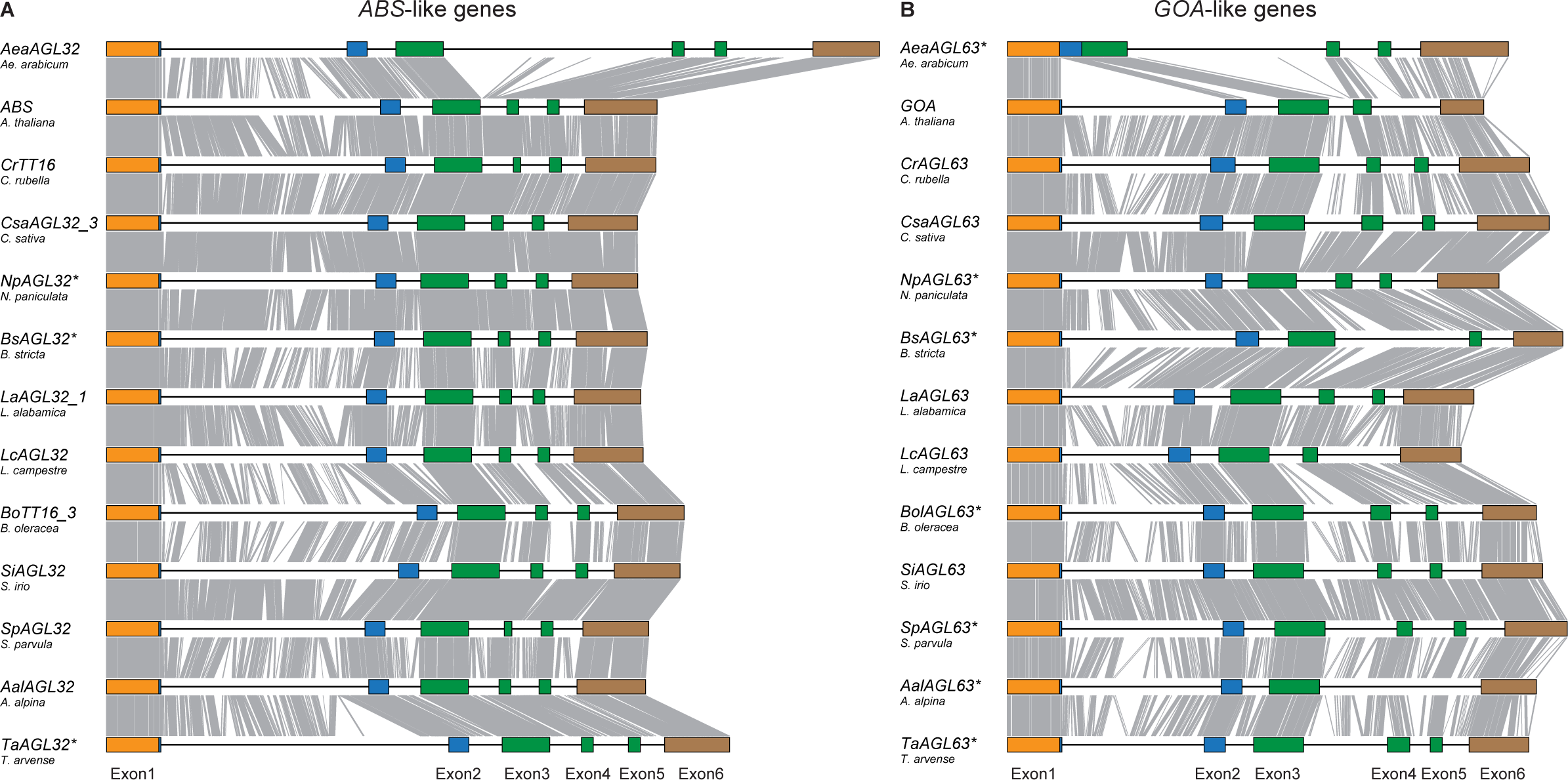
Exon-intron-structure alignment of *ABS*- (A) and *GOA*-like (B) genes. Exons are shown as boxes, introns as horizontal lines. Colors indicate domains encoded by the exons as follows: orange, MADS domain; blue, I domain; green, K domain; brown, C domain. Vertical lines indicate identical nucleotides in neighboring genes in a multiple sequence alignment. A star next to the gene name indicates that the exon-intron structure is based on a prediction solely using genomic sequence. Exon numbers are given at the bottom.

### Sequence changes in GOA-like proteins

To investigate which consequences the accelerated evolution of the nucleotide sequences of *GOA*-like genes have had on the protein level, we analyzed alignments of ABS- and GOA-like proteins (Supplemental Figure 5). We found that in Brassicaceae the average pairwise protein sequence identity of GOA-like proteins (61%) is significantly lower than that of ABS-like proteins (85%) (Mann–Whitney U-Test, p<0.001) (Supplemental Table 2), which is compatible with less purifying selection on coding regions of *GOA*-like genes and the exon-intron structure analyses presented above.

Comparing ABS- and GOA-like proteins, we observe a number of amino acid differences in the highly conserved MADS domain, which is responsible for DNA binding (Supplemental Figure 6). The respective changes increase the number of positively charged amino acids in GOA-like proteins, which in turn significantly increases the pI values of these proteins (Mann–Whitney U-Test, p<0.001) (Supplemental Figure 6). Since the DNA backbone is negatively charged we hypothesized that GOA-like proteins bind stronger to DNA than ABS-like and other MADS-domain proteins. Therefore, we determined the thermodynamic dissociation constant Kd of binding of ABS, GOA and, for comparison, SEP3 (a class E floral homeotic protein) of *A. thaliana* to a *cis*-regulatory element (CArG-box) known to bind MADS-domain proteins well (Jetha et al., 2014) (Supplemental Figure 7). Unexpectedly, we found that GOA binds less strongly to the DNA-sequence element (K_d_ = 3.50 ± 1.07 nM) than ABS (K_d_ = 0.37 ± 0.14 nM) or SEP3 (K_d_ = 1.78 ± 0.52 nM). We hypothesize that other amino acid substitutions in the MADS domain (Supplemental Figure 6), such as the substitution of the generally highly conserved arginine by lysine at position 3 (Melzer et al., 2006), or changes in the protein conformation, might counteract the increase of the pI values.

We also detected great differences between ABS- and GOA-like proteins in the K domain encoded by exons 3, 4, and 5. From the crystal structure of SEP3 it is known that the amino acids encoded by exons 3 and 4 mainly form a dimerization interface, whereas tetramerization is mostly achieved by the amino acids encoded by exon 5 (Puranik et al., 2014). The GOA-like proteins of *L. campestre* and *A. thaliana* have shortened K domains (Supplemental Figure 5), as the sequence homologous to exon 5 is not present in the mRNA due to mutations at the 3’ splice site and the branching point, respectively, of intron 4 (numbering according to Figure 6). Furthermore, shortened K domains are predicted for the GOA-like proteins of *Boechera stricta* and *Arabis alpina* (Supplemental Figure 5). For the species in which the mRNA encoding for the GOA-like protein contains all exons coding for the K domain, we observe differences in the probability to form coiled-coils as compared to the K domains of ABS-like proteins. For ABS-like proteins two regions with high probabilities of coiled-coil formation are predicted (Figure 7). For the analyzed GOA-like proteins (all containing all exons encoding for the K domain) the beginning of the two putative α-helices is shifted towards the C terminus by several amino acids and most intriguingly, the coiled-coil probability for helix 2 is quite low. It is thus quite likely that only one helix is formed, which, however, is shortened in comparison to ABS-like proteins. The inability to form a second helix is likely due to insertions and substitutions in the protein part encoded by exon 4 (Supplemental Figure 5). These insertions and substitutions disrupt the sequential pattern of hydrophobic amino acids necessary for the formation of a second helix.

**Figure 7:**
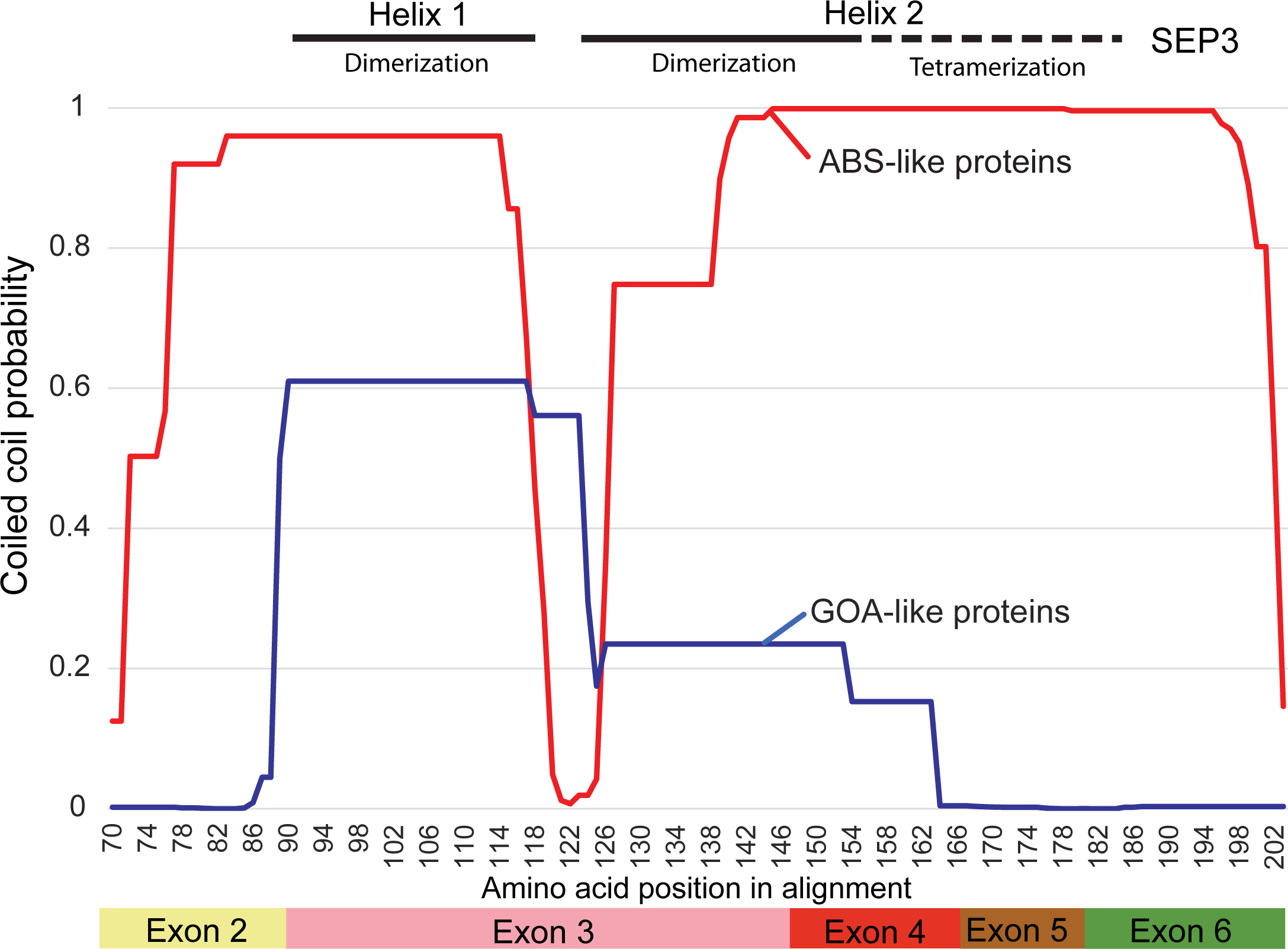
Coiled-coil formation probability for ABS- and GOA-like proteins of Brassicaceae. Coiled-coils prediction was performed based on the alignment in Supplemental Figure 5. Shown is only the prediction for a part of the alignment. Boxes at the bottom indicate by which exons this part is encoded. Positions orthologous to the positions of SEP3 forming two α-helices responsible for dimerization as well as tetramerization (Puranik et al., 2014), are indicated above the diagram as black lines.

Also in the C-terminal domain, which is involved in transcription activation and higher order complex formation in some MIKC-type proteins (Honma and Goto, 2001; Lange et al., 2013), all analyzed Brassicaceae GOA-like proteins differ dramatically from the other eudicot B_sister_ proteins (Supplemental Figure 5). Firstly, these proteins lack the derived PI motif at the C terminus and secondly, exon 6 is prolonged at the 5’ end in several species most likely due to changes in splice sites.

### ABS-like proteins show a conserved pattern of protein-protein interactions

ABS from *A. thaliana* has been shown to interact with the MIKC-type MADS-domain proteins SEP1, SEP2, SEP3, SEP4 and AP1 as well as with the non MIKC-type MADS-domain proteins AGL74, AGL97 and AGL92 in Yeast Two-Hybrid (Y2H) assays (de Folter et al., 2005; Kaufmann et al., 2005). To test whether the interactions with MIKC-type proteins are also conserved for ABS-like proteins from other Brassicaceae, we performed a Y2H analysis (Supplemental Table 3). All tested ABS-like proteins were able to form heterodimers with AP1 and all ABS-like proteins except CrTT16 from *Capsella rubella* interacted with SEP1, SEP3 and SEP4 from *A. thaliana*. For SEP2, we found that BhAGL32 from *Boechera holboellii* forms stable heterodimers. AalAGL32 from *A. alpina* and CrTT16 form weak heterodimers with SEP2, whereas ABS from *A. thaliana*, LcAGL32 from *L. campestre* and ThAGL32 from *Tarenaya hassleriana* were unable to do so in our assay (Supplemental Table 3). For GOA only one interaction partner, AGL16, was identified previously by a direct search against other MIKC-type proteins (de Folter et al., 2005), and could be confirmed by us (Supplemental Table 4). We tried to identify other possible interaction partners by screening a cDNA library from *A. thaliana* covering all plant organs at several developmental stages. In two independent experiments we were unable to reproducibly identify interaction partners. Furthermore, we were unable to affirm the interaction with AGL16 found by Y2H analysis in BiFC experiments (Supplemental Figure 8). Moreover, the GOA orthologs of *C. rubella* and *L. campestre* are unable to form heterodimers with the *A. thaliana* AGL16 protein (Supplemental Table 4). Thus, while ABS-like proteins show a number of conserved interaction partners, our findings show that GOA has no or only a very limited number of interaction partners, at least in *A. thaliana*.

### In Brassicaceae transcript accumulation of *GOA*-like genes is much lower than that of *ABS*-like genes

We then compared the expression level of *ABS* and *GOA* from *A. thaliana* to several orthologs from Brassicaceae using quantitative reverse transcription PCR (qRT-PCR) (Figure 8). Very low expression was observed in roots and leaves for all analyzed genes. *ABS* was found to be expressed most strongly in young siliques, followed by flowers and floral buds (Figure 8 A). *GOA* expression was generally found to be very low; highest expression levels were found in floral buds, but equaling only 8% of the *ABS* transcript abundance. Expression was also observed in roots, flowers, and siliques but at an even lower level than in floral buds. In *C. rubella*, the expression pattern observed for *CrTT16* was different to that of its *A. thaliana* ortholog *ABS* (Figure 8 B). We found *CrTT16* expression to be restricted to floral buds, flowers, and siliques. In contrast to *ABS*, the expression level of *CrTT16* decreased in siliques. The expression of the *GOA*-like gene *CrAGL63* was detected to be very low in floral buds and flowers and almost no transcripts were observed in roots, leaves, and siliques. The expression pattern of the *ABS* orthologs *AalAGL32* (from *A. alpina*) and *EsAGL32* (from *E. salsugineum*) was found to be similar to the expression pattern of *ABS* (Figure 8 C, D) with transcript accumulation being highest in young fruits. For the *GOA* ortholog of *A. alpina (AalAGL63)* we detected a very low level of transcript abundance in floral buds and flowers, and no expression in roots, leafs and siliques.

**Figure 8:**
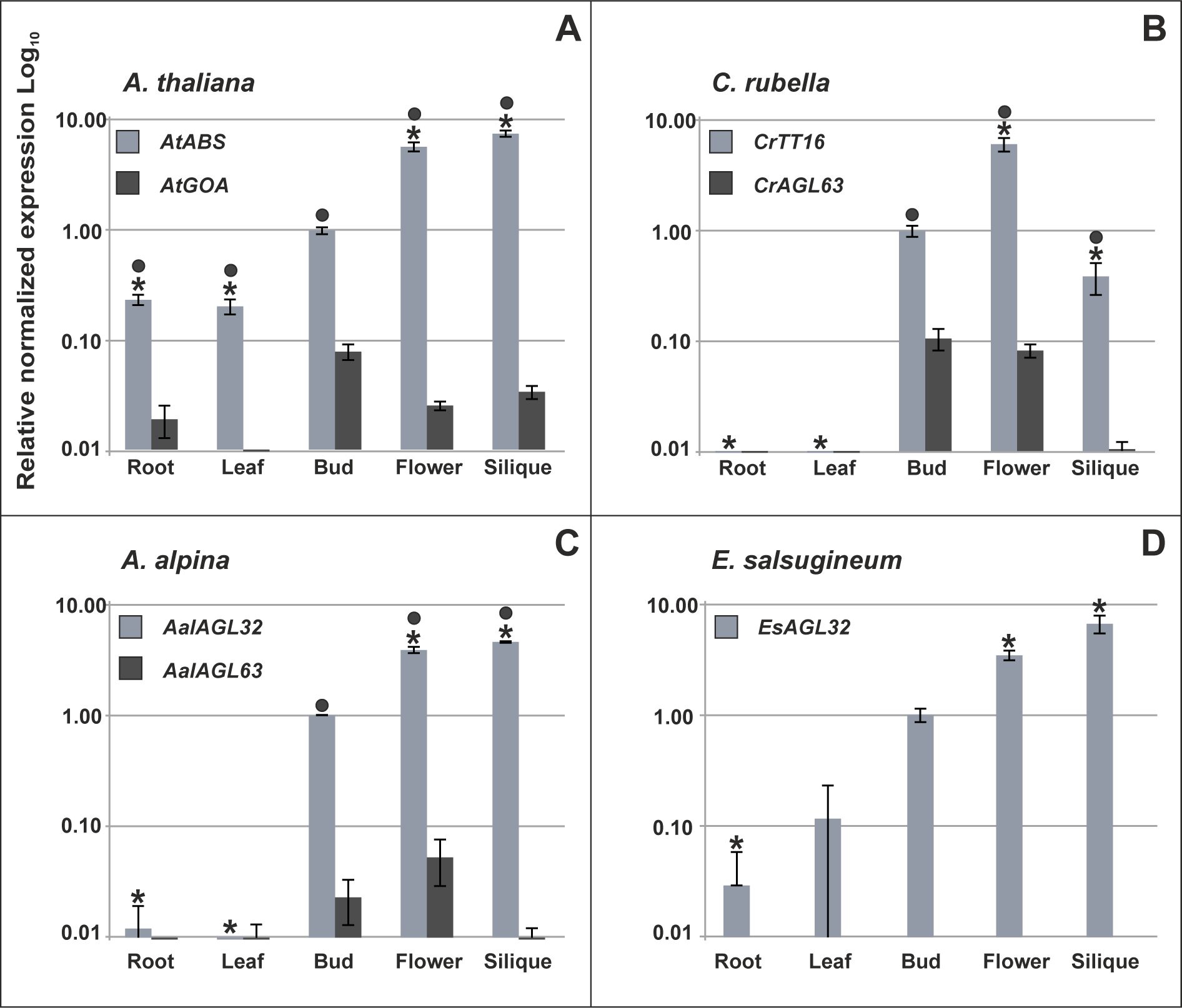
Quantitative real-time-PCR expression analysis of *ABS*- and *GOA-like genes from selected Brassicaceae*. Expression quantities of mRNAs in roots, leaves, flowers and siliques are presented relative to the expression level of the corresponding *ABS*-like gene in floral buds, which was set to 1. Bars represent standard deviations. Relative expression levels of **(A)** *ABS* and *GOA* from *A. thaliana*. **(B)** *CrTT16* (*ABS* ortholog) and *CrAGL63 (GOA* ortholog) from *C. rubella* **(C)** *AalAGL32* (*ABS* ortholog) and *AalAGL63* (*GOA* ortholog) from *A. alpina* and **(D)** *EsAGL32* (*ABS* ortholog) from *E. salsugineum*. A star indicates a statistically significant (P value <0.05) difference between the expression of an *ABS-*like gene in different tissues as compared to its expression in floral buds and a circle indicates a significant (P value <0.05) difference between the expression of *ABS*- and *GOA*-like genes in the same tissue. Note that the expression levels are shown on logarithmic (Log_10_) scale and that there is no *GOA*-like gene in *E. salsugineum*

In summary, except for *CrTT16* all analyzed *ABS*-like genes were expressed in a consistent manner in various members of the Brassicaceae, with the highest expression in young fruits followed by expression in flowers and floral buds. Remarkably, expression levels of *GOA*-like genes were always determined to be one to three orders of magnitude lower than those of *ABS*-like genes.

### The *A. thaliana goa-1* mutant does not show a mutant fruit phenotype

Our findings suggest that, in contrast to the highly conserved *ABS*-like genes, many *GOA*-like genes of Brassicaceae have been functionally compromised, nonfunctionalized or have even been lost. However, for *GOA* from *A. thaliana* a novel function in fruit expansion was previously reported using constitutive overexpression (Erdmann et al., 2010) and mutant analysis (Prasad et al., 2010). To determine as to whether *GOA* has indeed been neofunctionalized, we reanalyzed the T-DNA mutant line *goa-1* investigated by Prasad et al. (2010), which is assumed to be a null mutant. Using PCR and sequencing, we first found that the T-DNA is not inserted in exon 3 as described by Prasad et al. (2010), but located within intron 4 (Supplemental Figure 9). However, we were not able to amplify a full length *goa* cDNA from the *goa-1* mutant line, suggesting that it is indeed a null mutant. Furthermore, we did not observe any obvious phenotypic differences between wildtype and *goa-1* plants regarding growth habit and appearance of leaves, flowers, and fruits under standard long-day conditions. Since a function in fruit growth was proposed for *GOA* (Prasad et al., 2010), we measured the fruit length, width and seed number per fruit in three independent experiments performed in two different labs with independently grown plants. However, we were not able to detect a significant difference between wildtype and *goa-1* fruits (Supplemental Table 5). Thus our data do not rule out that even *GOA* of *A. thaliana* is a pseudogene.

## DISCUSSION

### A morbid interest in gene death

Genes are somewhat like organisms: they are ‘born’, they ‘live’ for a while, and eventually they ‘die’ and get lost, involving processes such as gene duplication, neo- or subfunctionalization, and nonfunctionalization. Nonfunctionalization is probably the most frequent fate of duplicated genes (Panchy et al. (2016) and references cited therein), but compared to neo- and subfunctionalization it has received much less scientific interest. Often only global, genome-wide parameters have been determined, such as the “propensity” or “rate” of gene loss (Krylov et al., 2003; Borenstein et al., 2007). One reason might be that gene loss is often considered to be just the trivial outcome of random mutations in sequences without a function on which purifying selection is not acting anymore. This general neglect of gene death is unfortunate since there is evidence that gene loss can be of considerable adaptive value (reviewed in Albalat and Cañestro (2016), examples from humans and birds provided by MacArthur et al. (2007); Daković et al. (2014)). Moreover, if gene death occurs slowly, e.g. under weak purifying selection, it is still possible that due to a mutation neofunctionalization kicks in before definitely deleterious mutations (such as large deletions) have occurred (Schilling et al., 2015). Thus in contrast to lost genes, slowly dying genes might represent a pool of DNA sequences for neofunctionalization and innovation.

Unfortunately, many dying genes, especially if lacking clear hallmarks of pseudogenes such as precocious stop codons or reading frame shifts, hamper genome annotation, simply because it is often not obvious as to whether these genes still function, do not function anymore or do function again. Moreover, genes with a high propensity of gene loss may lead to erroneous conclusions in approaches that use the absence of specific genes as characters to reconstruct organismic relationships. It has been argued that gene loss data harbor some of the most robust phylogenetic information (Sharma et al., 2014; Schierwater et al., 2016). However, our case study on *GOA*-like genes argues for caution. If some established genes in some clades of organisms are prone to undergo nonfunctionalization and thus get lost relatively frequently in an independent, convergent way, the absence of such a gene in other taxa may erroneously suggest close relationships of taxa lacking the gene even though they are not closely related at all.

In conclusion, the tempo and mode of gene loss deserve more scientific interest. Here we report an especially intriguing case of a whole gene clade that erodes in a convergent way in a well-defined family of angiosperms (Brassicaceae) after having survived for many millions of years in the lineage that led to Brassicaceae. The case is particularly striking because it affects a clade of MIKC-type MADS-box genes. These genes are generally well-known for their strong conservation during angiosperm evolution (Gramzow and Theißen, 2015). Studying the degeneration of a clade of MIKC-type genes at unprecedented detail provides novel insights into the life and death of genes and gene families.

### On necessity and chance: evolution of *ABS*- and *GOA*-like genes outside Brassicaceae

For a deeper understanding of gene death a reasonably well understanding of gene birth is essential. It was previously suggested that Bsister genes duplicated into *ABS*- and *GOA*-like genes in the eurosid II clade in the lineage that led to extant Brassicaceae (Erdmann et al., 2010). Our phylogeny reconstructions suggest that the respective gene duplication event occurred before the divergence of asterids and rosids (Figure 1; Supplemental Figures 1 and 2), probably by the triplication of the ancestral Bsister gene during the γ-WGT of the common ancestor of asterids and rosids 117 to 123 MYA (Jiao et al., 2012; Vekemans et al., 2012; Magallón et al., 2015).

Analyzing the fate of the two clades of Bsister genes after duplication reveals that quite a number of eudicot lineages have lost either the *GOA*-like gene or the *ABS*-like gene, but never both (Figures 1 and 3). This observation is easily explained by the requirement of at least one copy of an important Bsister gene but an arbitrary loss of the other copy, even though alternative scenarios cannot be excluded. Our data are compatible with the view that it did not matter in many lineages as to whether the *ABS*- or *GOA*-like gene maintained a canonical B_sister_ gene function, so that the other copy could get lost, suggesting that both copies were initially functionally redundant. The similarity of the expression patterns of the ‘surviving’ *ABS*- or *GOA*-like gene (summarized in Supplemental Figure 3) corroborates this hypothesis.

Interestingly, however, some plant lineages have maintained both *ABS*- and *GOA*-like genes until today, i.e. for roughly 120 million years. Extant plants with both kinds of genes include *V. vinifera*, *P. trichocarpa*, both *Citrus species* analyzed as well as many Brassicaceae species (Figures 1 and 3, Supplemental Figure 1 and 2). Since data about the function of *GOA*-like genes outside of Brassicaceae are essentially absent, and gene expression data are scarce (Supplemental Figure 3), it is impossible to say as to whether maintenance of both gene types was dominated by subfunctionalization, neofunctionalization, or a combination of both (subneofunctionalization). Even more interesting, we also identified some lineages which had kept both *ABS*- and *GOA*-like genes for many millions of years, but then still lost either the one or the other gene in an apparently stochastic way (Figure 1 and 3; Supplemental Figure 1 and 2). For example, the *ABS*-like gene was lost in the lineage that led to *C. papaya*, after the lineage that led to *T. hassleriana* (Cleomaceae) and Brassicaceae had branched-off roughly about 70 to 80 MYA (Bell et al., 2010; Magallón et al., 2015). This finding implies that both genes had been retained for around 40 to 50 million years before the *ABS*-like gene was lost. Conversely, the *GOA*-like gene was lost in the lineage that led to *T. hassleriana* after the lineage that led to extant Brassicaceae had branched off from a common ancestor around 40 MYA (Magallón et al., 2015). In this case, the *GOA*-like gene was retained for roughly 80 million years in the respective lineage before it got lost. All in all, we identified 5 cases each of loss of *ABS-* and *GOA*-like genes outside of Brassicaceae (Figure 3). This is a minimal, conservative estimate of the total loss of genes because a denser taxon sampling would likely uncover many more cases.

The pattern of apparently arbitrary loss of either one or the other gene may suggest functional redundancy of both genes for a remarkably long time. We consider complete redundancy unlikely, however, since it should lead to nonfunctionalization and a consequent loss of one copy within around 17 million years, as found for *A. thaliana* (Lynch and Conery, 2003). On the other hand, scenarios of classical sub-or neofunctionalization may also not apply here, since these would be difficult to reconcile with a pattern of arbitrary loss of either *ABS*- or *GOA*-like genes; both neo- and classical subfunctionalization should lead to a long-term conservation of both kinds of genes. We argue that ‘dosage subfunctionalization’ may explain the pattern of evolution of *ABS*- and *GOA*-like genes to a large extent. Analyzing gene duplicates using whole genome data from yeast (Qian et al., 2010) and several *Paramecium* species (Gout and Lynch, 2015) revealed a retention of ‘apparently’ redundant genes for a long time. The authors proposed a model according to which the expression of both gene copies is required after gene duplication to obtain a dosage of the gene product high enough to fulfill the ancestral function (Qian et al., 2010; Gout and Lynch, 2015). Eventually, random genetic drift may lead to an uneven expression level of the two duplicates long after duplication. This way, one of the two copies may acquire such a low expression level by random mutations that it only contributes in an (almost) insignificant way to the overall gene function. Then, further deleterious mutations may accumulate or the gene may get lost by random genetic drift, even after many millions of years of co-existence with its paralog (Qian et al., 2010; Gout and Lynch, 2015).

Since little is known about the dynamics of gene loss in plants, and especially dosage subfunctionalization has not been documented in detail in this branch of life yet, we decided to investigate the evolution of B_sister_ genes in more detail in Brassicaceae due to the unprecedented genomics resources available for this plant family.

### Delayed convergent degeneration of *GOA*-like genes in Brassicaceae

In the lineage that leads to Brassicaceae both *ABS*- and *GOA*-like genes have been retained for a remarkable period of (roughly) 80 million years (Figure 3). Then, however, convergent gene death only affecting *GOA*-like genes set in within Brassicaceae (Figure 4). We thus describe a novel pattern of duplicate gene evolution, Delayed Convergent Asymmetric Degeneration (DCAD), affecting only one clade of a pair of paralogous clades after dozens of millions of years of co-existence (Figure 9). Since both neo- and classical subfunctionalization should have led to a long-term conservation of *GOA*-like genes also in Brassicaceae (Innan and Kondrashov, 2010), we argue that the DCAD phenomenon requires an alternative explanation and that dosage subfunctionalization is involved.

**Figure 9:**
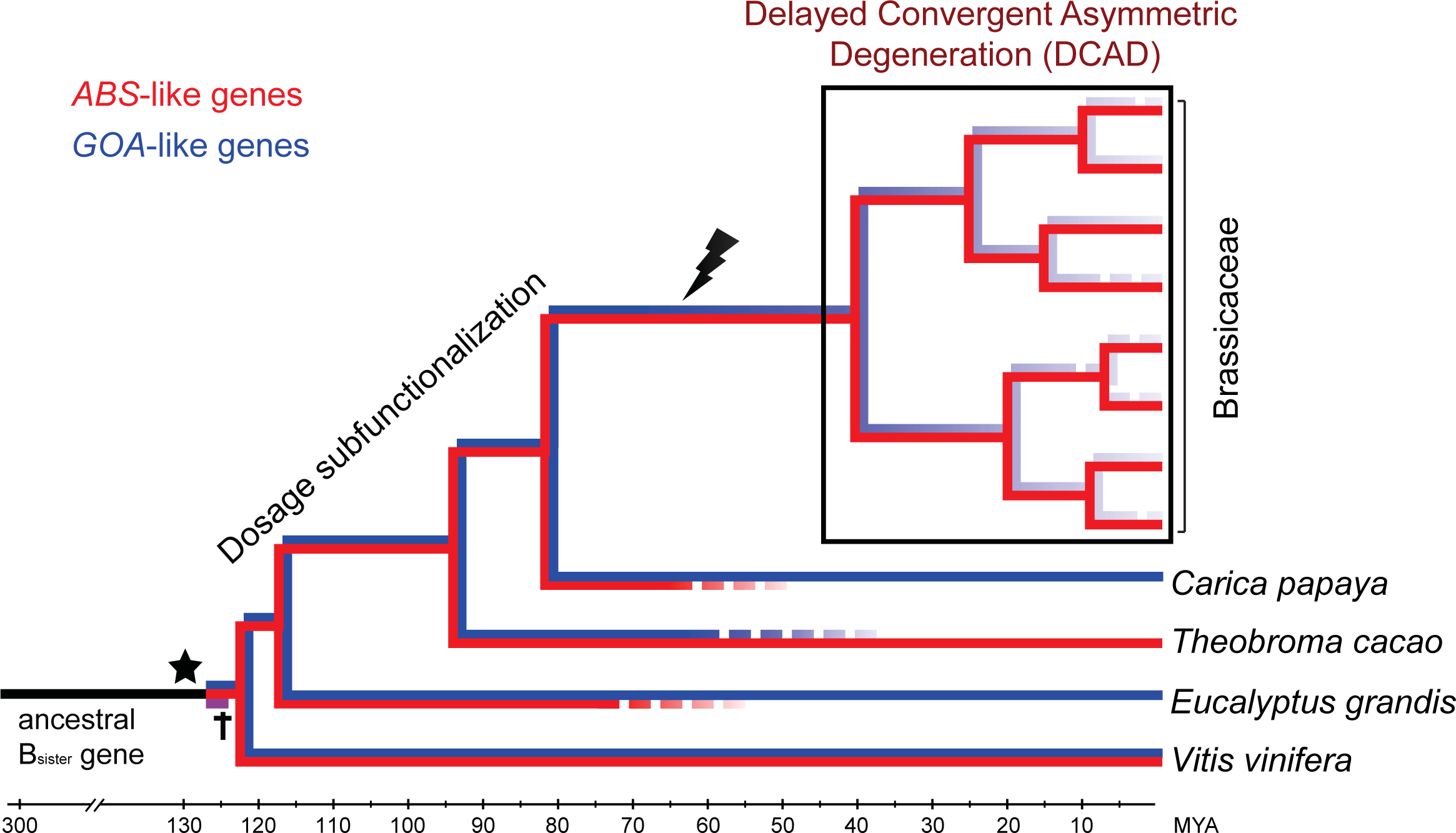
The scenario of Delayed Convergent Asymmetric Degeneration (DCAD) during paralogous gene evolution. DCAD is exemplified with *ABS*-like genes (shown in red; strongly conserved) and *GOA*-like genes (shown in blue; undergoing DCAD). In an initial phase, two paralogous genes may co-exist for many million years, e.g. due to dosage subfunctionalization. DCAD sets in when in the stem group of a taxon (here: Brassicaceae) one of the paralogs (here: the *GOA*-like gene) catches a mutation (lightning symbol) that compromises its function seriously (but maybe not completely), affecting most (if not all) orthologous genes in the crown group of that taxon (boxed). The triplication of the ancestral B_sister_ gene is indicated by a star, the putatively lost third paralog is labeled with a black cross. Rough age estimates are given in Million Years Ago (MYA) at the bottom.

We find that most of the analyzed *GOA*-like genes are somewhat compromised and probably impaired in their function due to deleterious mutations, are *en route* to being lost, or have even already been lost in several cases (Figure 4). However, we identified many different mutations distinguishing the *GOA*-like genes of Brassicaceae from other B_sister_ genes, ranging from single nucleotide polymorphisms (SNPs) to larger indel mutations. The observed mutations essentially affect all parts of MIKC-type genes, including introns (Figure 6). Mutations in exons lead to different changes in the amino acid sequences of GOA-like proteins, including substitutions in the highly conserved MADS domain (Supplemental Figures 5, 6) and indels in the K domain (Supplemental Figure 5); the latter apparently compromised the ability of the K domains of GOA-like proteins from Brassicaceae to form coiled-coils (Figure 7) and thus their ability to interact with other MIKC-type proteins. Indeed, we were not able to identify another interaction partner for GOA from *A. thaliana* in our Y2H experiments apart from AGL16 (Supplemental Table 4), whose interaction could moreover not be confirmed by the BiFC experiments (Supplemental Figure 8). In contrast we confirmed the known interactions with E-class proteins (SEP1 to SEP4) (Kaufmann et al., 2005; de Folter et al., 2006) for the analyzed Brassicaceae ABS-like proteins (Supplemental Table 3), which have a classical K domain (Figure 7).

The diversity of mutational changes observed in *GOA*-like genes of Brassicaceae suggests that these mutations do not represent the initial mutational causes of the convergent degeneration of *GOA*-like genes in Brassicaceae, but may have been ‘passengers’ rather than ‘drivers’ of these degeneration processes. In line with this notion we find elevated ω (dN/dS) values for *GOA*-like genes in Brassicaceae species, indicating reduced selective pressure. The ω values of *GOA*-like genes in Brassicaceae are much higher than those observed for most other MIKC-type MADS-box genes (Purugganan et al., 1995; Mondragon-Palomino et al., 2009; Lee and Irish, 2011), including B_sister_ genes outside of Brassicaceae for which we found an ω value of 0.20 (Figure 5). The value of 0.46 for *GOA*-like genes in Brassicaceae is in between the value for canonical MADS-box genes and the average value for pseudogenes (ω = 0.59) as identified in *A. thaliana* (Zou et al., 2009; Yang et al., 2011), corroborating the view that *GOA*-like genes have lost functional importance during the evolution of Brassicaceae.

Since *GOA*-like genes in diverse Brassicaceae are affected in a convergent way (Figure 4) it seems reasonable to assume that an initial mutation determining the fate of these genes likely happened in the stem group of extant Brassicaceae after the lineage that led to Caricaceae had already branched-off. This raises the question as to which initial event doomed *GOA*-like genes of Brassicaceae to extinction. All *GOA*-like genes of Brassicaceae that have been investigated were found to be expressed at extremely low levels in floral buds and flowers, and often at even lower levels in siliques, where high expression of a classical B_sister_ gene is expected (Bernardi et al., 2014) (Figure 8). A common denominator of all *GOA*-like genes in Brassicaceae thus seems to be their much lower expression level as compared to *ABS*-like genes, especially in developing fruits (siliques). It is thus conceivable that an initial mutation in a regulatory region, such as the promoter, of an ancestral *GOA*-like gene in the stem group of Brassicaceae significantly lowered the expression level of this gene diminishing its functional importance in comparison to the ancestral *ABS*-like gene in a dosage-subfunctionalization scenario.

The reduced expression, and consequently lowered functional relevance and reduced selection pressure on *GOA*-like genes during the radiation of crown group Brassicaceae may then have allowed for the accumulation of the different additional putative ‘passenger’ mutations. These mutations may have reduced the functionality of these genes even more, and eventually may have led in many cases to complete pseudogenization or even gene loss.

Our analyses did not reveal any case of complete loss of *ABS*-like genes in any species of Brassicaceae. Moreover, we found that the expression pattern of *ABS*-like genes from the analyzed Brassicaceae species is consistent with the previously identified expression pattern of other B_sister_ genes (Becker et al., 2002; de Folter et al., 2006; Yang et al., 2012; Chen et al., 2013; Lovisetto et al., 2013). Since *ABS*-like genes are also strongly conserved in terms of function as revealed by mutant phenotypes, where available (Deng et al., 2012; Mizzotti et al., 2012), it appears reasonable to assume that these genes largely maintain the ancestral function of B_sister_ genes in ovule and seed development within Brassicaceae whenever the *GOA*-like paralog is too much hampered in its function.

Our observations on the substitution rates and expression patterns of *GOA*-like genes in Brassicaceae certainly matches findings of genome-wide studies of different organisms: the probability of a gene to get lost is proportional to the ω value below 1 and inversely proportional to the strength of the gene’s expression and the number of its protein interaction partners (Krylov et al., 2003; Borenstein et al., 2007).

As genes approaching ω values of 1 show very low expression and (as far as known) few or no protein interaction partners, *GOA*-like genes of Brassicaceae may indeed fit the bill of a whole subclade of genes in the state of dying. This ultimately leads to the question as to whether there is any functional *GOA*-like gene left in Brassicaceae. For most genes, only proxies of gene function are available. These typically do not provide clear hallmarks of any functional relevance. However, *AalAGL63* from *A. alpina*, for example, is expressed just above the detection limit of qRT-PCR (Figure 8C) and a large part of the K-domain encoded by exons 4 and 5 in other GOA-like proteins is missing in AalAGL63 (Supplemental Figure 5), hence the protein likely cannot form dimers with any of the *A. alpina* SEP proteins. We hypothesize that most, if not all, of the genes with characteristics similar to that of *AalAGL63* might be pseudogenes.

The only *GOA*-like gene from a Brassicaceae for which a mutant phenotype has ever been reported is *GOA* from *A. thaliana*. In this case, neofunctionalization that led to a function in fruit growth was reported (Erdmann et al., 2010; Prasad et al., 2010). However, our analysis of the same T-DNA insertion line of *GOA* from *A. thaliana* which was previously reported to reveal the mutant knock-out phenotype (Prasad et al., 2010) did not reveal a mutant fruit phenotype under different environmental conditions. Thus, based on our data, even *GOA* from *A. thaliana* might well be a pseudogene. Recently, however, (Xu et al., 2016) reported that *ABS* (*TT16*) and *GOA* redundantly promote nucellus degeneration during seed development in *A. thaliana*. This function may explain why *GOA* does not show neutral evolution (as would be revealed by an ω value of about 1). However, redundancy is not easy to reconcile with the considerable differences between *ABS* and *GOA* in terms of structure and expression level and pattern, and would not be evolutionarily stable. Like the role of *GOA* in fruit development, also its role in nucellus degeneration may thus deserve further investigations.

### DCAD beyond GOA

Candidates for DCAD might be recognized by careful phylogeny reconstruction involving, if available, complete sets of gene family members from all relevant species, including also all genes that are expressed at an extremely low level only (thus requiring genomics data). An indication of DCAD would be phylogenetic gene trees where one of two long existing paralogs has representatives in (almost) all relevant species, whereas the sister clade appears to lack orthologs in many species, suggesting convergent, asymmetric (i.e. biased) gene loss after quite some time of coexistence. This way, the remaining sister clade members may be identified as putative pseudogenes even though they may not yet show clear hallmarks of pseudogenization. Thus a DCAD pattern of gene evolution might be developed into heuristics that aid in gene and genome annotations. More detailed studies may then support or reject the DCAD hypothesis and the pseudogene identity of individual sequences.

We thus think that DCAD is not a unique pattern of *GOA* vs. *ABS* gene evolution. We rather argue that it represents a more general scenario of duplicate gene evolution that may have been observed before, but that has not been defined, fully recognized and appreciated as such. Importantly, DCAD may be more frequent than one may assume.

DCAD may have indeed shaped the molecular evolution of a considerable number and diversity of genes. For example, *OsMADS30*-like genes of grasses (Poaceae) represent another case of genes from the Bsister family of MADS-box genes. In contrast to their closely related and highly conserved *OsMADS29*-like paralogs, *OsMADS30*-like genes appear to evolve in a DCAD-like pattern, even though indications of complete pseudogenization are less obvious (Schilling et al., 2015). Also, other MIKC-type MADS-box genes of grasses show a DCAD pattern of gene evolution, such as *AP1/FUL*-like (also termed *SQUA*-like) genes; in this case, a subclade termed *FUL4*-like genes, appears to degenerate in a DCAD-like way, whereas the other three subclades are quite conserved (Wu et al., 2017). The *PISTILLATA*- (*PI*-) like gene of legumes may also evolve in a DCAD-like pattern. Whereas e.g. *MtPI* of *Medicago truncatula* seems to have maintained the ancestral role of a typical class B floral organ identity gene specifying petals and stamens, its paralog *MtNGL9* is expressed much weaker, evolved faster, does not reveal any mutant floral phenotype after loss-of-function, and might well be on a way towards pseudogenization – despite coexistence with *MtPI* for about 50 to 60 million years (Roque et al., 2016). *CAULIFLOWER* vs. *APETALA1* from *A. thaliana* may represent another kind of DCAD-like evolution, disfavoring *CAL* by the loss of several transcription factor binding sites and favoring *AP1* by gaining these (Ye et al., 2016).

The DCAD pattern of gene evolution can also be found in gene families encoding transcription factors other than MADS-domain proteins. For example, a pattern similar to the one we describe here as DCAD has been reported for the *DICHOTOMA* (*DICH*) vs. *CYCLOIDEA* (*CYC*) genes of the TCP family of Antirrhineae (Hileman and Baum, 2003). However, due to a lack of genome sequences, it was difficult to conclusively demonstrate the absence of *DICH* genes from the genomes of species from which they could not be isolated. In the discussion of Hileman and Baum (2003) on why *DICH* genes, in contrast to *CYC* genes, are not all maintained but slowly degenerate and get lost, they come close to our model of DCAD based on dosage subfunctionalization. It includes the hypothesis that *DICH* genes may have gotten lost independently once they crossed the threshold to functional insignificance, e.g. due to reduced purifying selection or reduced expression, beyond which purifying selection could prevent gene loss. This hypothesis may well also apply to the *GOA* scenario.

Also, genes encoding general (basal) transcription factors may reveal a DCAD-like pattern of molecular evolution. An example concerns *TFIIAγ*, which encodes a small subunit of TFIIA that is required for RNA polymerase II function. Two paralogous genes, *TFIIAγ1* and *TFIIAγ5*, originated in the stem group of grasses (Poaceae) roughly about 50 to 80 MYA and have both been maintained e.g. in several extant rice (*Oryza*) species (Sun and Ge, 2010). In this case, *TFIIAγ5* appears to have maintained the ancestral function as a subunit of the TFIIA complex, whereas *TFIIAγ1* displays many features of the DCAD syndrome, including accelerated evolution, relaxed purifying selection, reduced expression and gene loss (Sun and Ge, 2010).

Our examples deliberately focus on gene families encoding transcription factors. We are confident, however, that readers which are more familiar with other types of genes, such as those encoding enzymes, will quite easily find cases of DCAD-like gene evolution also in their favorite gene families. Once appreciated as a general scenario of duplicate gene evolution, DCAD patterns are not difficult to find.

## MATERIALS AND METHODS

### Isolation and amplification of genomic DNA

We cloned genomic sequences of both the *ABS* and *GOA* orthologs of *L. campestre* and *A. alpina*; of the *ABS* orthologs of *E. salsugineum* and *T. hassleriana*; and of the *GOA* orthologs of *A. lyrata* and *B. rapa*. Detailed information for plant growth can be found in the Supplemental Methods. Genomic DNA of *L. campestre* was isolated using the NucleoSpin^®^ Plant II DNA kit (Macherey&Nagel). Genomic DNA of the different *B. rapa* subspecies and *A. lyrata* was isolated according to Edwards et al. (1991). Genomic DNA of *T. hassleriana, A. alpina* and *E. salsugineum* was extracted using the DNeasy^®^ Plant Mini kit (Qiagen). The amplification was done using Phusion^®^ polymerase (Thermo Scientific and New England BioLabs). All primer sequences can be found in the Supplemental Data Set 2.

### RNA isolation and cDNA preparation

For the following species, we obtained cDNA of both the *ABS* and *GOA* orthologs experimentally: *A. alpina*, *C. rubella*, *L. campestre* and *S. irio;* and of *ABS* orthologs of *Ae. arabicum, B. holboellii*, *E. salsugineum*, *S. parvula*, and *T. hassleriana*. Detailed information for plant growth can be found in the Supplemental Methods. Plant material was collected and immediately processed for RNA extraction or snap frozen in liquid nitrogen and stored at −80°C. Total RNA from *L. campestre, Ae. arabicum* and *S. irio* were isolated using the Plant-RNA-OLS kit (OMNI Life Science). Total RNA from *B*. *holboellii* was extracted using TRItidy G™ (AppliChem). Total RNA from *T. hassleriana, A. alpina, C. rubella* and *E. salsugineum* was extracted using the NucleoSpin^®^ RNA Plant kit (Macherey&Nagel). cDNA synthesis was performed with the RevertAid H Minus Reverse Transcriptase (Thermo Scientific) using 500 ng to 3 μg of total RNA and primer AB05 for the rapid amplification of cDNA ends (RACE). Random hexamer primers were used for cDNA production for the qRT-PCR. cDNA was produced using an oligo(dT) primer for the amplification of the coding sequences from cDNA. Subsequently, the cDNA was amplified using gene-specific primers. A 3’-RACE for the *ABS*- and *GOA*-like genes of *L. campestre* was performed using 2 μl of cDNA purified by the “NucleoSpin Gel and PCR Clean-up Kit” (Macherey&Nagel) with gene-specific primers and primer AB07 using Phusion polymerase (Thermo Scientific). A 5’-RACE was performed as follows: 10 μl purified cDNA was poly(A)-tailed with 1.5 μl terminal deoxynucleotidyl transferase (Thermo Scientific). 5 μl of the poly(A)-tailed cDNA was amplified using primer AB05 and a gene-specific primer with Phusion polymerase. A “nested” PCR was performed with Phusion polymerase using 5 μl of the first PCR product, primer AB07, and a gene-specific nested primer. PCR products were cloned and sequenced. All primer sequences can be found in Supplemental Data Set 2.

### Phylogeny reconstruction

Sequences of B_sister_ proteins as identified previously from eudicots (Becker et al., 2002; Leseberg et al., 2006; Díaz-Riquelme et al., 2009), monocots (Yang et al., 2012; Gramzow and Theißen, 2013), basal angiosperms (Becker et al., 2002) and gymnosperms (Becker et al., 2003; Carlsbecker et al., 2013) or within this work were used. Detailed information for the *in silico* identification of B_sister_ genes can be found in the Supplemental Methods. APETALA3, a B class protein from *A. thaliana* was added to the dataset. All sequences were aligned using Probalign (Roshan and Livesay, 2006). The alignment was cropped using trimAL (Capella-Gutiérrez et al., 2009) with the parameters -gt 0.9 and -st 0.0001. The phylogenies were reconstructed using RAxML (Stamatakis, 2006) and MrBayes (Ronquist and Huelsenbeck, 2003). For RAxML, the protein substitution model LG (Le and Gascuel, 2008) was chosen, and the maximum likelihood phylogeny and 1000 bootstrap replicates were calculated in one program run (option “-f a”). For MrBayes, the protein substitution model WAG (Whelan and Goldman, 2001) was chosen, six million generations were calculated sampling every 100th generation and excluding the first 25% generations from the consensus tree reconstruction containing all compatible groups (contype=allcompat). The phylogenies were rooted using APETALA3 as representative of the outgroup.

### Analysis of exon-intron structures of *ABS*-and *GOA*-like genes from Brassicaceae

Exon-intron structures of *ABS*- and *GOA*-like genes of Brassicaceae were determined by an alignment of cDNA to genomic sequences. However, for the *GOA*-like genes of *Ae. arabicum*, *A. alpina*, *S. parvula* and *B. oleracea*, we had to rely on gene predictions because despite extensive efforts we were not able to amplify cDNA from these genes, possibly due to extremely low levels of expression (see, e.g. Figure 8). Furthermore, for the *ABS*- and *GOA*-like genes of *Neslia paniculata, B. stricta*, and *Thlaspi arvense* we also relied on gene predictions. The genomic sequences were taken from the translation start to the stop codon. To have the same number of genes in each dataset, one gene was chosen arbitrarily from *C. sativa*, *L. alabamica* and *B. oleracea*.Genome sequences were aligned for *ABS*- and *GOA*-like genes separately using MLAGAN (option: “Use translated anchoring in LAGAN/Shuffle-LAGAN”) (Brudno et al., 2003). The graphical presentation of the exon-intron structures including the alignment was produced using a customized perl script (Supplemental Methods). For calculation of pairwise identity percentages, the tool SIAS (Sequences Identities and Similarities server, http://imed.med.ucm.es/Tools/sias.html) was used with standard parameters. The alignment was split into exons and introns and the method PID3 was chosen. For the values obtained, the average was calculated for each exon and intron separately.

### Selection analysis

We conducted a selection analysis for *ABS*- and *GOA*-like genes, in which we considered only genes of species which have retained both clades of B_sister_ genes. The B_sister_ gene of *Aquilegia coerulea* was used as representative of the outgroup. The same number of genes was used for *ABS*- and *GOA*-like genes. Therefore, the same genes that were chosen for the exon-intron analysis for *C. sativa*, *L. alabamica* and *B. oleracea* were used. Nucleotide sequences were translated into protein sequences using EMBOSS Transeq (Rice et al., 2000), aligned using Probalign (Roshan and Livesay, 2006), manually checked using Jalview (Waterhouse et al., 2009), and back-translated using RevTrans (Wernersson and Pedersen, 2003). The nucleotide alignment was cropped using trimAL (Capella-Gutiérrez et al., 2009) with the parameter -gt 0.8. Corrected distances were plotted versus uncorrected distances of transitions and transversions as calculated in PAUP* 4.0beta10 (Wilgenbusch and Swofford, 2003) to exclude saturation of nucleotide substitutions. The nucleotide alignment and a phylogeny of these genes based on a species phylogeny (Koenig and Weigel, 2015) were used as input for codeml of the PAML 4.5 package (Yang, 1997; Piwarzyk et al., 2007). We used two branch models, a one-ratio (M0) (Anisimova et al., 2001) and a five-ratio model. The five ratio model allows one ω ratio for all the branches in the clade of Brassicaceae *GOA*-like genes; one ω ratio for the branch leading to the *GOA*-like genes of Brassicaceae; one ω ratio for all the branches in the clade of Brassicaceae *ABS*-like genes; one ω ratio for the branch leading to the *ABS*-like genes of Brassicaceae; and one ω ratio for all remaining branches. The models were compared using a likelihood ratio test (LRT).

### Protein sequence analysis

For the determination of pairwise protein sequence identity, we aligned sequences of B_sister_ proteins from basal eudicots, ABS and GOA-like proteins from species where both types of proteins are present, as well as the sequence of the GOA-like protein from *C. papaya* using Probalign (Roshan and Livesay, 2006), and calculated the corresponding values using the SIAS server (http://imed.med.ucm.es/Tools/sias.html). The values for the isoelectric points (pI) of the MADS domains were obtained using the program ProtParam (Gasteiger et al., 2005). For the coiled-coils analysis, the sequences of ABS- and GOA-like proteins were aligned using ProbCons (Do et al., 2005). Single amino acids wrongly placed at exon borders were corrected manually (Supplemental Figure 5). Subalignments of ABS- and GOA-like proteins from Brassicaceae were obtained by deleting the corresponding other sequences from the alignment and were then taken as input for the program PCOILS (Lupas, 1996) (window size 28 amino acids). For creating Figure 7, the positions of the results from the coiled-coiled prediction were mapped according to the positions of the original alignment.

### Quantitative Reverse Transcription PCR (qRT-PCR)

The qRT-PCR assay for the expression analysis of *ABS-* and *GOA*-like genes in different plants was performed as described in Chen et al. (2013). Amplification efficiencies for each gene were calculated using a cDNA dilution series and also by linear regression calculations for every reaction using LinRegPCR 2013.0 (Ruijter et al., 2009). Standard dose response curves were performed for all the genes analyzed. Each reaction was performed in biological and technical triplicates along with water and RNA controls for each primer pair. For *A. alpina*, *E. salsugineum* and *C. rubella*, the expression was normalized against *ACTIN7* (*ACT7*) and *GAPC1.* For *ABS* and *GOA* from *A. thaliana*, expression was normalized against *PEROXIN4* (*PEX4*) and *AT4G33380*. These genes were chosen from Czechowski et al. (2005) to discriminate subtle differences in expression patterns of genes with very low expression. The data was processed by the relative standard curve method and the fold difference between the expression of internal controls and the genes of interest calculated using the comparative Cq method (ΔΔCq) (Livak and Schmittgen, 2001). To compare the expression levels of *ABS*- and *GOA*-like genes in each species, the expression level of the *ABS*-like gene was set to 1 for the value in floral buds. A one-way ANOVA was performed to calculate the statistical significance of the differences between the expression values. Oligonucleotide sequences are provided in Supplemental Data Set 2.

### Yeast 2-Hybrid and BiFC assays

The Y2H assays for protein interaction were performed as described previously (Lange et al., 2013). All constructs were analyzed for auto-activation and unspecific DNA binding. All Y2H experiments were performed in three biological replicates. The full-length coding sequence of *GOA* was cloned into pGBKT7 (Clontech, San Jose, CA, USA) and used as bait to screen against an *A. thaliana* cDNA library provided by Hans Sommer (Max-Planck-Institute for Plant Breeding Research, Cologne, Germany). The screen was performed according to Kaufmann et al. (2005). The BiFC experiments were performed according to Lange et al. (2013). For the assay, the coding sequences (CDS) of *GOA* and *AGL16* were amplified from pGADT7/GOA and pGADT/AGL16 constructs (Erdmann et al., 2010), respectively, without their native stop codon, and cloned into pNBV-YC and pNBV-YN vectors (Walter et al., 2004).

### In vitro transcription/translation and EMSA

The cDNA of *GOA* and *ABS* from *A. thaliana* were amplified by PCR and cloned into pTNT (Promega) using *EcoRI* and *SmaI* recognition sites. The plasmid pTNT_SEP3 was used from Melzer et al. (2009). *In vitro* translation was done using the T7 TNT Quick Coupled Transcription/Translation mix (Promega). To determine the Kd values for the DNA binding of SEP3, ABS and GOA dimers, a DNA probe containing a consensus CArG-box sequence derived from the regulatory intron of *AGAMOUS* was used in EMSAs (Jetha et al., 2014). In these assays, a constant amount of protein (2 μl of the *in vitro* translation solution containing the protein) was titrated against increasing concentrations of DNA (1.31, 2.62, 5.23, 10.47, 20.94, 41.87, 52.34, and 62.81 nM in each reaction, respectively) as described in Jetha et al. (2014).

### Analysis of the *A. thaliana goa-1* mutant

The *goa-1* mutant (SALK_061729C) described by Prasad et al. (2010) as well as the *A. thaliana* Col-0 wildtype were obtained from the Nottingham Arabidopsis Stock Centre (NASC, http://arabidopsis.info). The plants were grown under long-day conditions as described in the plant growth section (Supplemental Methods). DNA was isolated from leaf material according to Kasajima et al. (2004). Plants were genotyped using the primers LP, RP and LBb1.3 (Supplemental Data Set 2). The length and width of 4 to 5 consecutive stage-17b siliques from the primary inflorescence were measured from 10 to 12 plants per genotype. For measurement of seed number, siliques were cleared prior to microscopy in a mixture of chloral hydrate/glycerol/water (8:2:1) overnight at 4°C. Microscopy was performed using a Leica M205 FA microscope system.

## Accession Numbers

Sequence data from this article can be found in Genbank under the accession numbers given in Supplemental Data Set 1.

## Supplemental Data

**Supplemental Data Set 1:** List of sequences used in this study.

**Supplemental Data Set 2:** Primer Sequences for the amplification of *ABS*-like, *GOA*-like, and B_sister_ genes or cDNA.

**Supplemental Figure 1:** Phylogeny of B_sister_ genes of seed plants as reconstructed by Bayesian inference based on protein sequences.

**Supplemental Figure 2:** Maximum Likelihood phylogeny of B_sister_ genes in seed plants reconstructed based on protein sequences.

**Supplemental Figure 3:** Expression pattern of *ABS*- and *GOA*-like genes of different eudicots.

**Supplemental Figure 4:** Alignment of the 5’ part of the MADS-box sequences of the GOA orthologs of *Brassica oleracea* and of different *Brassica rapa* subspecies.

**Supplemental Figure 5:** Alignment of B_sister_ proteins of basal eudicots (BS*) and ABS- and GOA-like proteins.

**Supplemental Figure 6:** Alignment of the MADS domains of B_sister_ proteins of basal eudicots (BS*) and of ABS- and GOA-like proteins.

**Supplemental Figure 7:** Analysis of binding of SEP3 (A, B), ABS (C, D) and GOA (D, E) to DNA by electrophoretic mobility shift assays (EMSAs).

**Supplemental Figure 8:** Protein dimerization analysis by bimolecular fluorescence complementation (BiFC).

**Supplemental Figure 9:** Location of the T-DNA insertion in the *GOA* gene of the *goa-1* mutant line (*A. thaliana*).

**Supplemental Table 1:** Phylogenetic Analysis by Maximum Likelihood (PAML) results and results of the Likelihood Ratio Test for selection analysis.

**Supplemental Table 2:** Pairwise amino acid identities in percent of B_sister_ proteins of basal eudicots and of ABS- and GOA-like proteins.

**Supplemental Table 3:** Summary of the protein interaction analysis of ABS-like proteins from Brassicales as revealed by Yeast Two-Hybrid analysis.

**Supplemental Table 4:** Summary of the protein interaction analysis of GOA-like proteins as revealed by Yeast Two-Hybrid analyses.

**Supplemental Table 5:** Silique measurements of *goa-1* and wildtype plants of stage-17b fruits.

**Supplemental Methods:** Plant growth and *in silico* identification of B_sister_ genes.

## ACKNOWLEDGEMENTS

We thank Tim Sharbel (Gatersleben, Germany), Dmitry Smetanin (Zürich, Switzerland), George Coupland (Cologne, Germany) and Dong-Ha Oh (Urbana-Champaign, IL, USA) for sequence information. We thank Hans Sommer (Cologne, Germany) for providing the *A. thaliana* cDNA library for Y2H analysis. Furthermore, we thank Julian Schenk (Gießen, Germany) for helping with *C. rubella* and *E. salsugineum* cloning and expression analysis, Kai Pfannebecker (Gießen, Germany) for one of the Y2H cDNA screens for *GOA* and Dietmar Haffer (Gießen, Germany) for growing the *B. holboellii* plants, Florian Rümpler (Jena, Germany) for help with statistical analysis and coiled-coils predictions and Ulrike Coja (Osnabrück, Germany) and Heidi Kressler (Jena, Germany) for excellent technical assistance. The work was supported by the German Research Foundation DFG, grant TH 417/9-1 to G. T. and BE 2547/9-1 to A. B.

## AUTHOR CONTRIBUTIONS

A.B. and G.T. designed the research, interpreted and evaluated data. A.H. performed the synteny analysis, analysis of gene loss, sequence analysis, structure predictions as well as the bioinformatic analysis of the expression pattern of B_sister_ genes and *goa-1* mutant analysis. L.G. performed the phylogenetic and selection analysis, exon-intron alignment, sequence searches as well as the bioinformatic analysis of the expression pattern of B_sister_ genes. N.K. performed sequence searches and analysis, synteny analysis and *goa-1* mutant analysis. A.G. performed the EMSA analysis. A. S.B. performed and analyzed the qRT-PCR experiments. O. S. performed the protein interaction assays. K.M. provided seed material, DNA and cDNA sequences as well as crucial information regarding Brassicaceae phylogeny. A.H., L.G. and G.T. wrote the manuscript.

